# The Wnt effector TCF7l2 promotes oligodendroglial differentiation by repressing autocrine BMP4-mediated signaling

**DOI:** 10.1101/2020.09.16.300624

**Authors:** Sheng Zhang, Yan Wang, Xiaoqing Zhu, Lanying Song, Xinhua Zhan, Edric Ma, Jennifer McDonough, Hui Fu, Franca Cambi, Judith Grinspan, Fuzheng Guo

## Abstract

Promoting oligodendrocyte differentiation represents a promising option for remyelination therapy for treating the demyelinating disease multiple sclerosis (MS). The Wnt effector TCF7l2 was upregulated in MS lesions and had been proposed to inhibit oligodendrocyte differentiation. Recent data suggest the opposite yet underlying mechanisms remain elusive. Here we unravel a previously unappreciated function of TCF7l2 in controlling autocrine bone morphogenetic protein (BMP4)-mediated signaling. Disrupting TCF7l2 results in oligodendroglial-specific BMP4 upregulation and canonical BMP4 signaling activation *in vivo*. Mechanistically, TCF7l2 binds to *Bmp4* gene regulatory element and directly represses its transcriptional activity. Functionally, enforced TCF7l2 expression promotes oligodendrocyte differentiation by reducing autocrine BMP4 secretion and dampening BMP4 signaling. Importantly, compound genetic disruption demonstrates that oligodendroglial-specific BMP4 deletion rescues arrested oligodendrocyte differentiation elicited by TCF7l2 disruption *in vivo*. Collectively, our study reveals a novel connection between TCF7l2 and BMP4 in oligodendroglial lineage and provides new insights into augmenting TCF7l2 for promoting remyelination in demyelinating disorders such as MS.

**Significance Statement:** Incomplete or failed myelin repairs, primarily resulting from the arrested differentiation of myelin-forming oligodendrocytes from oligodendroglial progenitor cells, is one of the major reasons for neurological progression in people affected by multiple sclerosis (MS). Using *in vitro* culture systems and *in vivo* animal models, this study unraveled a previously unrecognized autocrine regulation of BMP4-mediated signaling by the Wnt effector TCF7l2. We showed for the first time that TCF7l2 promotes oligodendroglial differentiation by repressing BMP4-mediated activity, which is dysregulated in MS lesions. Our study suggests that elevating TCF7l2 expression may be possible in overcoming arrested oligodendroglial differentiation as observed in MS patients.

## Introduction

Blocked differentiation of myelin-forming oligodendrocytes (OLs) from oligodendrocyte precursor (or progenitor) cells (OPCs) is a major cause for remyelination failure in multiple sclerosis, the most common demyelinating disorder (Franklin and Gallo, 2014). Promoting OL differentiation represents a promising option for remyelination therapy in combination with current immunomodulatory therapy for treating MS (Harlow et al., 2015). Identifying upregulated molecular targets in MS will provide new insights into designing remyelination therapeutics. The upregulation of the Wnt effector transcription factor 7-like 2 (TCF7l2) in MS brain lesions (Fancy et al., 2009) suggest that it might be a potential target for overcoming OL differentiation blockade (Fu et al., 2009).

TCF7l2 is one of the four members (TCF1, TCF7l1, TCF7l2, and LEF1) of the TCF/LEF1 family that transcriptionally activate the canonical Wnt/β-catenin signaling (Cadigan, 2012). Based on the inhibitory role of Wnt/β-catenin activation in OL differentiation (Guo et al., 2015), TCF7l2 had been previously proposed as a negative regulator of OL differentiation by transcriptionally activating Wnt/β-catenin signaling (Fancy et al., 2009; He et al., 2007). However, subsequent genetic studies have convincingly demonstrated that TCF7l2 is a positive regulator of OL differentiation that TCF7l2 play a minor role in activating Wnt/β-catenin in oligodendroglial lineage cells (Fu et al., 2009; Hammond et al., 2015; Zhao et al., 2016). In this context, the molecular mechanisms underlying TCF7l2-regulated OL differentiation remains incompletely understood. We previously reported that genetically disrupting TCF7l2 inhibited OL differentiation with concomitant upregulation of the canonical BMP4 signaling in the CNS (Hammond et al., 2015), suggesting that the Wnt effector TCF7l2 may promote OL differentiation by tightly controlling BMP4 signaling, an inhibitory pathway of OL development (Grinspan, 2020). In this study, we unraveled a new mechanism of TCF7l2 in promoting OL differentiation by repressing autocrine BMP4 signaling.

## Materials and Methods

### Animals

The transgenic mice used in our study were: *Olig2-CreER^T2^* (Takebayashi et al., 2002), *Plp-CreER^T2^* (RRID: IMSR_JAX:005975), Pdgfrα*-CreER^T2^* (RRID: IMSR_ JAX:018280), *Tcf7l2-floxed* (RRID: IMSR_JAX:031436), *Bmp4*-floxed (RRID: IMSR_JAX: 016878, the exon4 was flanked by two loxP sites). Cre transgene was always maintained as heterozygosity. All animals were from C57BL/6 background and maintained in 12 h light/dark cycle with water and food. Animal protocols were approved by the Institutional Animal Care and Use Committee at the University of California, Davis.

### Gene conditional KO by tamoxifen treatment

Tamoxifen (T5648, sigma) was dissolved in a mixture of ethanol and sunflower seed oil (1:9, v/v) at 30 mg/ml. All animals including cKO and control mice were intraperitoneally administered at a dose of 200 μg/g body weight.

### Magnetic-activated cell sorting (MACS)

The forebrain was dissociated by the papain dissociation kit (LK003176, Worthington) and dissociator (Miltenyi Biotec, #130-092-235).

Astrocyte and microglia were purified by anti-ACSA-2 micro beads (130-097-679, Miltenyi Biotec) and anti-CD11b micro beads (130-049-601, Miltenyi Biotec), respectively. The cell suspension was then incubated with anti-O4 micro beads (130-094-543, Miltenyi Biotec) for OPC purification.

### Rodent primary culture of oligodendrocyte precursor cells (OPCs) and differentiation

Primary mouse OPCs were isolated from cortices of pups from P0 to P2 using immunopanning procedure as described previously (Hammond et al., 2015; Zhang et al., 2020; Zhang et al., 2018b). Cerebral cortices were enzymatically digested using papain (20 U/ml, #LK003176, Worthington) supplemented with DNase I (250 U/ml; #D5025, Sigma) and D-(+)-glucose (0.36%; #0188 AMRESCO) and mechanically triturated to obtain single cell suspension. The mixed glial cells were plated on poly-D-lysine (PDL, #A003-E, Millipore) coated 10 cm dishes and cultured in DMEM medium (#1196092, Thermo Fisher) with 10% heat-inactivated fetal bovine serum (#12306-C, Sigma) and penicillin/streptomycin (P/S, #15140122, Thermo Fisher). After 24 hours incubation, cells were washed using HBSS (with calcium and magnesium, #24020117, Thermo Fisher), and cultured in serum-free growth medium (GM), containing 30% of B104 neuroblastoma medium and 70% of N1 medium (DMEM with 5 μg/ml insulin (#I6634, Sigma), 50 μg/ml apo-transferrin (#T2036, Sigma), 100 μM putrescine (#P5780, Sigma), 30 nM Sodium selenite (#S5261, Sigma), 20 nM progesterone (#P0130, Sigma)) until 80% of cellular confluency. The cells were then dissociated in single cell suspension and seeded to Thy1.2 (CD90.2) antibody (#105302, Biolegend) coated Petri-dish to deplete astrocytes, neurons and meningeal cells followed by incubation on NG2 antibody (#AB5320, Millipore) coated Petri-dish to select OPCs. OPCs were then cultured on PDL-coated plates with GM plus 5 ng/ml FGF (#450-33, Peprotech), 4 ng/ml PDGF-AA (#315-17, Peprotech), 50 μM Forskolin (#6652995, Peprotech,) and glutamax (#35050, Thermo Fisher). OPCs were differentiated in the differentiation medium, consisting of F12/high-glucose DMEM (#11330032, Thermo Fisher Scientific) plus 12.5 μg/ml insulin, 100 μM Putrescine, 24 nM Sodium selenite, 10 nM Progesterone, 10 ng/ml Biotin, 50 μg/ml Transferrin (#T8158, Sigma), 30 ng/ml 3,3′,5-Triiodo-L-thyronine (#T5516, Sigma), 40 ng/ml L-Thyroxine (#T0397, Sigma-Aldrich), glutamax and P/S.

### Vector-mediated TCF7l2 overexpression in primary oligodendrocytes and Oli-Neu cells

The expression vectors pFlag-CMV2-TCF7l2-E2, pFlag-CMV2-TCF7l2-S2, and pFlag-CMV2-TCF7l2-M1 were provided by Dr. Andreas Hecht at the University of Freiburg, Germany and were described in the original publication (Weise et al., 2010). For vector-mediated TCF7l2 overexpression in primary OPCs or Oli-Neu cells, the transfection was conducted using FuGENE6 kit (Cat#2619, Promega) according to the manufacturer’s protocol.

### Enzyme-linked immunosorbent assay (ELISA)

The medium collected from primary culture were used for BMP4 measurement assay at day 2 of differentiation. The BMP4 concentration was determined using the BMP4 ELISA kit (Cat#OKEH02960, Aviva System Biology). ELISA was performed according to our previous protocol (Zhang et al., 2020).

### Chromatin immunoprecipitation (ChIP)

ChIP assays were performed using a SimpleChIP Enzymatic Chromatin IP kit (#9003; Cell Signaling Technologies). Briefly, cells were cross-linked using 37% formaldehyde at a final concentration of 1% for 10 min at room temperature and then quenched with 125 mM glycine for 5 min. Cells were washed twice with cold PBS, harvested in PBS with protease inhibitor cocktail (PIC) and lysed to get nuclei. Chromatin was harvested, fragmented using micrococcal nuclease to length of approximately150-900 bp and then briefly sonicated to lyse nuclear membranes. The digested chromatin (500 μg) was subjected to immunoprecipitation with specific TCF7L2 antibody (Celling Signaling) or anti-Rabbit IgG (#2729 Cell signaling) as negative control overnight at 4° C with rotation and followed by ChIP-Grade protein G magnetic beads incubated for 2 hours at 4° C with rotation. After immunoprecipitation, the chromatin was eluted from the antibody/protein G magnetic beads and cross-links were reversed. DNA was purified and analyzed by RT-qPCR. Primer sequences (Integrated DNA Technologies) for the *Bmp4* binding site were as follows: forward primer 5′-GGTACCTGCACTTAAGCTTTGTCGG 3′, and Reverse primer: 5′-TCGTAGTCGCTGCACGCAG-3′. The data were quantified by percent input and performed in triplicates.

### Protein extraction and Western blot assay

Protein extraction and Western blot assay were performed as previously described with modifications (Wang et al., 2016; Zhang et al., 2020). Tissue or cells were lysed using N-PER™ Neuronal Protein Extraction Reagent (#87792, Thermo Fisher) supplemented with protease inhibitor cocktail (#5871S, Cell Signaling Technology) and PMSF (#8553S, Cell Signaling Technology) on ice for 10 min and centrifuged at 10,000 × g for 10 min at 4°C to pellet the cell debris. Supernatant was collected for concentration assay using BCA protein assay kit (#23225, Thermo Fisher Scientific). Protein lysates (30 μg) were separated by AnykD Mini-PROTEAN TGX precast gels (#456-9035, BIO-RAD) or 7.5% Mini-PROTEAN TGX precast gels (#456-8024, BIO-RAD). The proteins were then transferred onto 0.2 μm nitrocellulose membrane (#1704158, BIO-RAD) using Trans-blot Turbo Transfer system (#1704150, BIO-RAD). The membranes were blocked with 5% BSA (#9998, Cell signaling) for 1 hour at room temperature and incubated with primary antibodies at overnight 4 °C, followed by suitable HRP-conjugated secondary antibodies. Proteins of interest were detected using Western Lightening Plus ECL (NEL103001EA, Perkin Elmer). NIH Image J was used to quantify protein levels by analyzing the scanned grey-scale films. Primary and secondary antibodies used were: TCF7l2 (1:1000, RRID:AB_2199816, #2569S, Cell Signaling Technology), MBP (1:1000, RRID:AB_2139899, #NB600-717, Novus), CNPase (1:1000, RRID:AB_2082474, #2986, Cell Signaling Technology), HDAC2 (1:1000, RRID:AB_10624871, #5113, Cell Signaling Technology), TLE3 (1:1000, RRID:AB_2203743, #11372-1-AP, Proteintech), β-actin (1:1000, RRID:AB_330288, #4967L, Cell Signaling Technology), GAPDH (1:1000, RRID:AB_561053, #2118L, Cell Signaling Technology), and HRP goat anti-rabbit (1:3000, RRID:AB_228341, #31460, Thermo Fisher Scientific), anti-mouse (1:3000, RRID:AB_228307, #31430, Thermo Fisher Scientific) or anti-rat (1:3000, RRID:AB_10694715, #7077, Cell Signaling Technology) secondary antibodies.

### Protein co-immunoprecipitation (Co-IP)

Co-IP was performed as previously described with some modifications (Lang et al., 2013; Wang et al., 2017). Proteins were extracted by Pierce IP Lysis/Wash Buffer (#87787, Thermo Fisher) and protein concentration was evaluated using BCA protein assay kit. 1 mg of proteins were incubated with TCF7l2 antibody (#2569S, Cell Signaling Technology) or normal rabbit IgG (#2729, Cell Signaling Technology) as negative control for immunoprecipitation overnight at 4°C. Pierce Protein A/G Magnetic Beads (#88802, Thermo Fisher) were washed three times with Pierce IP Lysis/Wash Buffer and added to antigen/antibody complex for 1 hour at room temperature. Beads were then washed twice with IP Lysis/Wash Buffer and once with purified water. The antigen/antibody complex was eluted from beads by boiling in Lane Marker Sample Buffer (#39001, Thermo Fisher) containing β-mercaptoethanol for SDS-PAGE and Western blotting.

### RNA extraction, cDNA preparation, and RT-qPCR assay

Total RNA was isolated by RNeasy Lipid Tissue Mini Kit (#74804, QIAGEN) with on-column DNase I digestion to remove genomic DNA using RNase-Free DNase Set (#79254, QIAGEN). The concentration of RNA was determined by Nanodrop 2000. cDNA was synthesized by Qiagen Omniscript RT Kit (#205111, Qiagen). RT-qPCR was performed by using QuantiTect SYBR^®^ Green PCR Kit (#204145, QIAGEN) on Agilent MP3005P thermocycler. The relative mRNA level of indicated genes was normalized to that of the internal control gene Hsp90 (Hsp90ab1), using the equation 2^(Ct(cycle threshold) of Hsp90 – Ct of indicated genes) according to our previous protocol (Hull et al., 2020). The gene expression levels in control groups were normalized to 1. The RT-qPCR primers used in this study were as followed:

*Sox10* (F: ACACCTTGGGACAC GGTTTTC, R: TAGGTCTTGTTCCTCGGCCAT),
*Enpp6* (F: CAGAGAGATTGTGA ACAGAGGC, R: CCGATCATCTGGTGGACCT),
*Mbp* (F: GGCGGTGACAGACT CCAAG, R: GAAGCTCGTCGGACTCTGAG),
*Plp* (F: GTTCCAGAGGCCAACATCAAG, R: CTTGTCGGGATGTCCTAGCC)
*Mog* (F: AGCTGCTT CCTCTCCCTTCTC, R: ACTAAAGCCCGGATGGGATA C),
*Myrf* (F: CAGACCCAGGTGCTACAC, R: TCCTGCTTGATCATTCCGTTC),
*Bmp4-exon4:* (F: TGATCACCTCAACTCAACCAAC, R: TCATCCAGGTACAACATGGAAA),
*Tcf7l2* (F: GGAGGAGAAGAACTCGGAAAA, R: ATCGGAGG AGCTGTTTTGATT),
*Axin2* (F: AACCTATGCCCGTTTCCTCTA, R: GAGTGTAAAGACTTGGTCCACC),
*Naked1* (F: CAGCTTGCTGCATACCATCTAT, R: GTTGAAAAGGACGCTCCTCTTA),
*Sp5* (F: GTACTTGCCATCGAGGTAG, R: GGCTCGGACTTTGGAATC),
*Gfap* (GTGTCAGAAGGCCACCTCAAG/ CGAGTCCTTAATGACCTCACCAT),
Slc1a2 (CAACGGAGGATATCAGTCTGC/ TGTTGGGAGTCAATGGTGTC),
*Id1* (F: CTCAGGATCATGAAGGTCGC, R: AGACTCCGAGTTCAGCTCCA),
*Id2* (F: ATGGAAATCCTGCAGCACGTC, R: TGGTTCTGTCCAGGTCTCT),
*Id3* (CGACCGAGGAGCCTCTTAG/ GCAGGATTTCCACCTGGCTA),
*Id4* (CAGTGCGATATGAACGACTGC/ GACTTTCTTGTTGGGCGGGAT),
*Hsp90* (F: AAACAAGGAGATTT TCCTCCGC, R: CCGTCAGGCTCTCATATCGAAT).

### cRNA probe preparation, mRNA in situ hybridization (ISH), and combined mRNA ISH and immunohistochemistry (ISH/IHC)

Probe preparation, mRNA in situ hybridization, and combination of ISH/IHC were performed by using the protocols in our previous studies (Hammond et al., 2015; Zhang et al., 2020). In brief, we used a PCR-based approach to amplify 907bp and 623bp fragments of *Bmp4* and *Mag* mRNA, respectively. The primer sets we used were: *Bmp4*-forward: 5′-CCTGCAGCGATCCAGTCT-3′ / *Bmp4*-reverse: 5′-GCCCAATCTCCACTCCCT-3′; *Mag*-forward: 5′-CAAGAGCCTCTACCTGGATCTG-3′ / *Mag*-reverse: 5′-AGGTTCCTGGAGGTACAAATGA-3′. The core sequence (GCGATTTAGGTGACACTATAG) recognized by SP6 RNA Polymerase was attached to the 5′ of the reverse primers. The core sequence (GCGTAATACGACTCACTATAGGG) recognized by T7 RNA Polymerase was attached to the 5′ of the forward primers. The resulting PCR products were purified and used as templates for in vitro transcription by employing DIG RNA Labeling Kit (SP6/T7) (Roche #11175025910). The resulting RNAs generated from SP6-mediated in vitro transcription were used as positive probes (cRNA, complementary to the target mRNAs) whereas those generated from T7-mediated in vitro transcription were used as negative probes (same sequence as the target mRNA). Frozen sections (12μm thick) were used for mRNA ISH. Hybridization was performed in a humidity chamber at 65°C for 12 - 18 hours, followed by a series of stringent washing by saline-sodium citrate (SSC) buffer at 65°C (2x SSC 20 min for 3 times and then 0.2x SSC 20 min for 3 times) and RNase digestion at 37°C (30 min) to eliminate non-specific binding and non-hybridized mRNAs. The DIG signals were visualized by NBT/BCIP method (Roche, #11697471001). Depending on the abundance of target mRNAs, the duration of NBT/BCIP incubation (at 4°C) ranged from 4 hours to overnight.

For combined ISH/IHC, frozen sections (12μm thick) were used. DIG-conjugated cRNA probe hybridization was performed according to the procedure described above. After a series post-hybridization SSC washing and RNase digestion, the sections were briefly rinsed in TS7.5 buffer (0.1M Tris-HCl, 0.15M NaCl, pH7.5) followed by incubation in 1% blocking reagent (prepared in TS7.5 buffer) (Roche #11096176001) for 1 hour at room temperature. After blocking, a cocktail of anti-Sox10 antibody (1:100, #AF2864, R&D Systems) and AP-conjugated anti-DIG antibody (Roche #11093274910) diluted in 1% blocking reagent was applied and incubated at 4°C for overnight. The sections were then washed in TNT buffer (TS7.5 + 0.05% Tween-20) 10 min for 3 times. The primary antibody of Sox10 was visualized by incubating Alexa Fluor 488-conjugated secondary antibody at room temperature for 1 hour. The DIG-conjugated mRNA signals were visualized by HNPP/Fast Red Fluorescent Detection set (Roche #11758888001) according to the kit instructions.

### Tissue preparation and immunohistochemistry (IHC)

Tissue preparation and IHC were conducted as previously described (Hammond et al., 2015; Zhang et al., 2020; Zhang et al., 2018b). Mice were perfused with ice-cold PBS (#BP399-20, Fisher Chemical), and post-fixed in fresh prepared 4% paraformaldehyde (PFA, #1570-S, Electron Microscopy Science, PA) at room temperature for 2 hours. Tissues were washed three times (15 min per each time) using ice-cold PBS and cryoprotected using 30% sucrose (#S5-3, Fisher Chemical) in PBS at 4°C overnight followed by embedding in OCT prior. Twelve microns thick sections were serially collected and stored in −80 °C. For immunohistochemistry, slice was air dry in room temperature for at least 1 hour, and blocked with 10% Donkey serum in 0.1% Triton X-100/PBS (v/v) for 1 hour at room temperature. Tissue was incubated with primary antibodies overnight at 4°C, washed three times in PBST (PBS with 0.1% Tween-20), and incubated with secondary antibodies at room temperature for 2 hours. DAPI was applied as nuclear counterstain. All images shown were obtained by Nikon C1 confocal microscope. An optical thickness of 10 μm was used for confocal z-stack imaging (step size 1 μm and 11 optical slices), and the maximal projection of the z-stack images was used for data quantification. Primary antibodies included Sox10 (1:100, RRID:AB_442208, #AF2864, R&D Systems), Sox10 (1:100, RRID:AB_2650603, #ab155279, Abcam), HDAC2 (1:100, RRID:AB_365273, #5113P, Cell Signaling Technology), TCF7l2 (1:100, RRID:AB_22199816, #2569S, Cell Signaling Technology), β-catenin (1:200, RRID:AB_476865, #7202, Sigma), P-Smad 1/5 (1:200, RRID:AB_491015, #9516, Cell Signaling Technology), CC1 (1:100, RRID:AB_2057371, #OP80, Millipore), PDFGR_α_ (1:100, RRID:AB_2236897, #AF1062, R&D Systems), APC(1:100, RRID:AB_2057493, sc-896, Santa Cruz Biotechnology), BLBP (1:100, RRID:AB_10000325, #ABN14, Millipore), Sox9 (1:200, RRID:AB_2194160,#AF3075, R&D system). All species-specific secondary antibodies (1:500) were obtained from Jackson ImmunoResearch.

### Immunocytochemistry (ICC)

Primary cells attached to the coverslips were fixed in 4% PFA for 15 min followed by rinse with PBS. The immunocytochemistry was performed as our previous studies (Guo et al., 2012; Hammond et al., 2015; Zhang et al., 2020). The following antibodies were used in our study: Sox10 (1:100, RRID: AB_2195374, #sc-17342,Santa Cruz Biotechnology; 1:500, RRID:AB_778021, #ab27655, Abcam), Mbp (1:500, RRID:AB_2564741, #SMI-99, Biolegend), TCF7l2 (1:200, RRID:AB_2199826, #2569S, Cell Signaling Technology), P-Smad1/5 (1:200, RRID:AB_491015, #9516, Cell Signaling Technology).

### Gene clone and luciferase activity assay

To generate a luciferase reporter under the control of the *intronic* promoter of the mouse *Bmp4* gene, a 439-bp DNA fragment containing the *Bmp4* promoter was amplified using the spinal cord genomic DNA with the forward primer 5′-<u>GGTACC</u>TGCACTTAAG***CTTTG***TCGG-3′ (underlined sequence is KpnI site, bold italic sequence is TCF7l2 binding site) and the reverse primer 5′-<u>CTCGAG</u>ACGGAATGGCTCCTAAAATG-3′ (underlined sequence is XhoI site). The PCR product was cloned into pGEM-T-Easy vector and confirmed by DNA sequencing. After digesting with KpnI and XhoI, the fragment was cloned into pGL2-Basic vector. The resulting luciferase reporter was designated as pGL2-Bmp4-Luc. Using pGL2-Bmp4-Luc as a template, TCF7l2-binding mutation constructs were generated by PCR using the above reverse primer and the following forward primer: <u>GGTACC</u>TGCACTTAAGTGGTGTCGG (underlined sequence is Kpn I site). The resulting mutated luciferase reporter was designated as pGL2-mBmp4-Luc. The dual luciferase assay was done in triplicate using TD-20/20Luminometer (Promega) according to the manufacturer’s instructions. Briefly, 0.25 μg of a luciferase reporter (Bmp4-Luc or mutBmp4-Luc), 0.25 μg of TCF7l2-S2 or TCF7l2-E2, and 5 ng of Renilla luciferase report (Promega) were co-transfected into U87 human primary glioblastma cells by using ESCORT V transfection reagent (Sigma). The fold increase in relative luciferase activity is a product of the luciferase activity induced by TCF7l2-S2 or TCF7l2-E2 (backbone vector pCMV) divided by that induced by an empty pCMV vector.

### Experimental designs and statistical analyses

All quantification was conducted by lab members blinded to genotypes or treatments. For in vivo gene cKO studies, animals were derived from multiple litters. Data were presented as mean ± standard error of mean (s.e. m.) throughout this manuscript. We used bar graph and scatter dot plots to show the distribution of the data. Each dot (circle, square, or triangle, if applicable) in the scatter dot plots represents one mouse or one independent experiment. We used Shapiro–Wilk approach for testing data normality. F test and Browne–Forsythe test were used to compare the equality of variances of two and three or more groups, respectively. The statistical analysis details of Figs. 1-3 were summarized in Table 1. P values of t test were displayed in each graph. Welch’s correction was used for Student’s t test if the variances of two groups were unequal. For comparisons among three or more groups with equal variances, ordinary one-way ANOVA was used followed by Tukey’s multiple comparisons, otherwise Welch’s ANOVA was used followed by unpaired t test with Welch’s correction. All data plotting and statistical analyses were performed using GraphPad Prism version 8.0. P value less than 0.05 was considered as significant, whereas greater than 0.05 was assigned as not significant (ns).

**Table 1 –.**
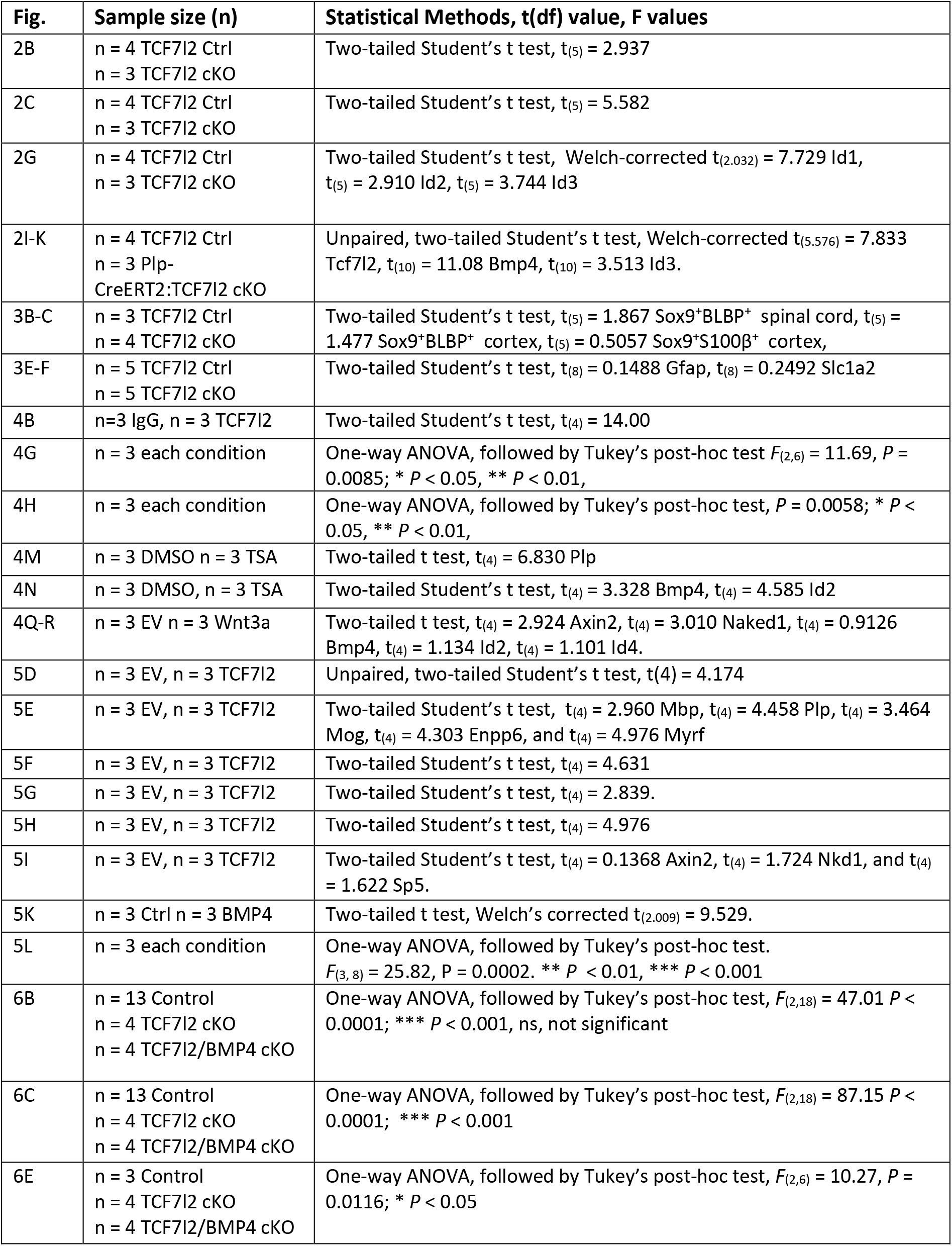
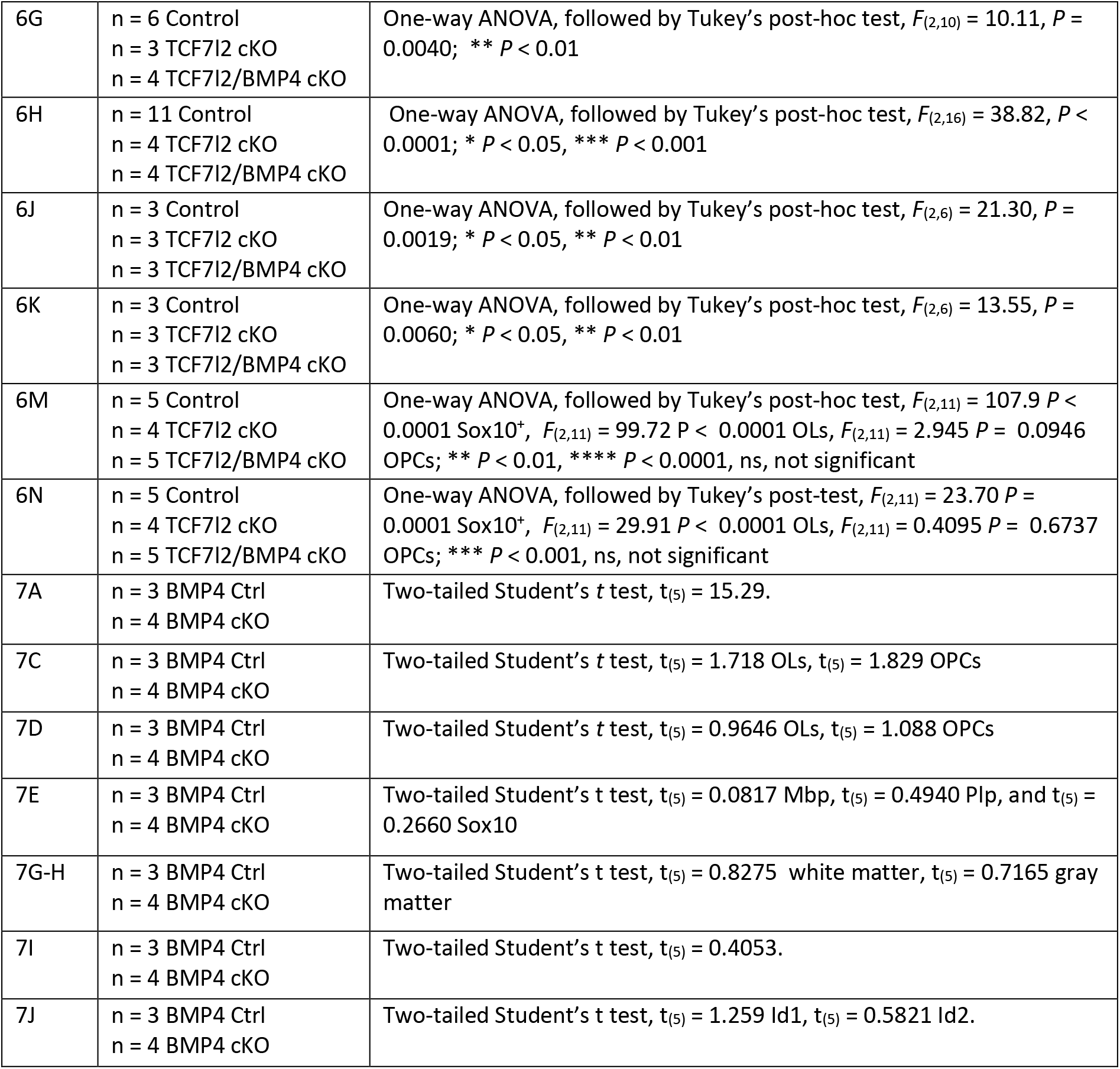
Detailed information of statistical analyses.

**Figure 1.**
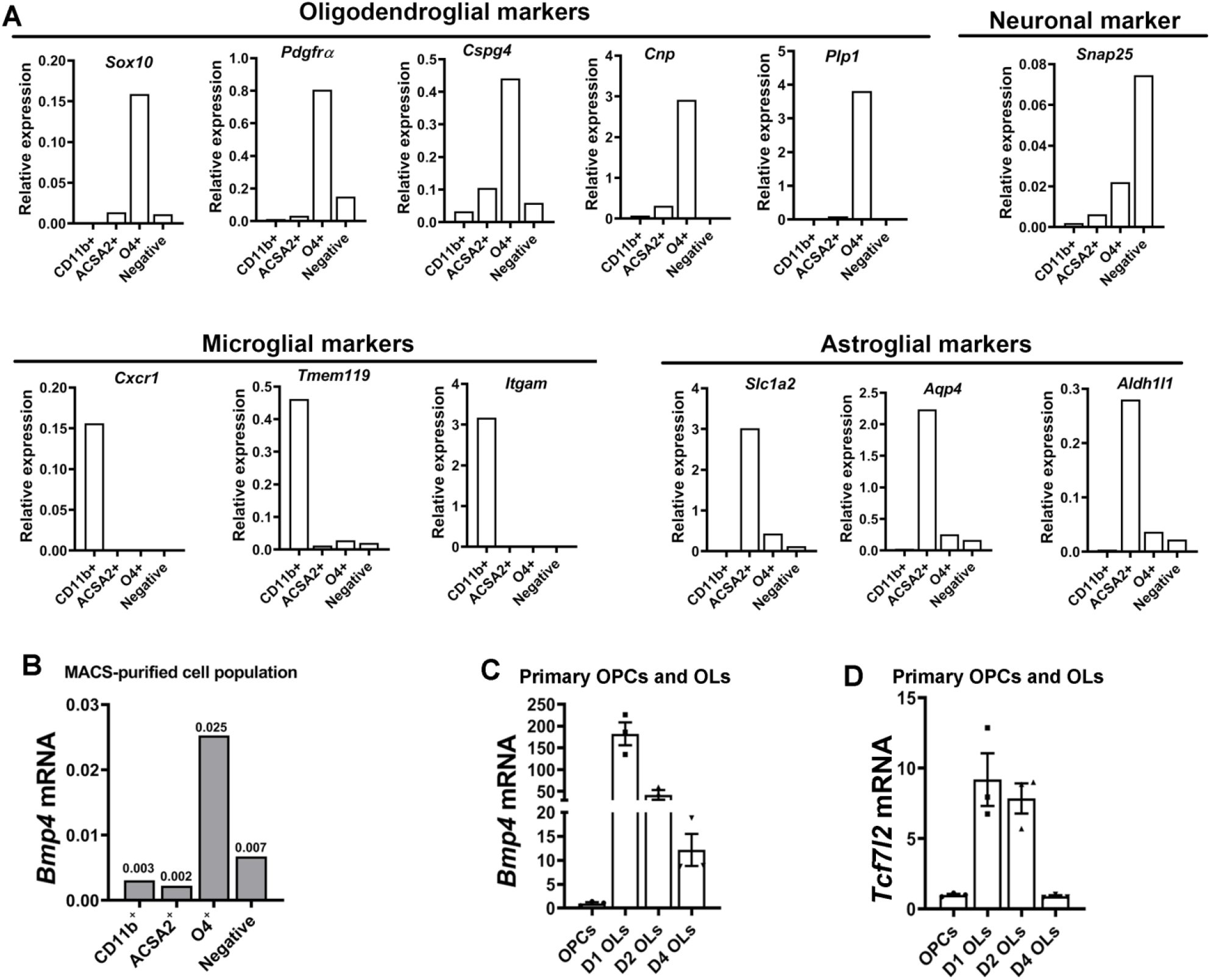
BMP4 is expressed primarily in newly differentiated oligodendrocytes. The brain of P5 C57BL/6 wild type mice (n=3, pooled together) was used for cell purification by magnetic-activated cell sort (MACS). CD11b, ACSA2 and O4-conjugated magnetic beads were used for positively selecting microglial, astroglial, and oligodendroglial lineage cells, respectively. Brain cells negative for CD11b, ACSA2, and O4 were designated as Negative components. RNA samples prepared for CD11b^+^, ASCA2^+^, O4^+^, and triple negative cell populations were used for RT-qPCR assays. **A**, RT-qPCR quantification validating the efficacy of MACS purification of each cell populations by lineage-specific markers (normalized to the internal control gene Hsp90 in each population). **B**, RT-qPCR assay for *Bmp4* in MACS-purified brain microglial (CD11b^+^), astroglial (ACSA2^+^), and oligodendroglial lineage (O4^+^) cells at P5. **C-D**, RT-qPCR assay for *Bmp4* (**C**) and *Tcf7l2* (**D**) in primary rodent OPCs maintained in proliferating medium and differentiating OLs maintained in differentiation medium for 1-4 days (D1-D4 OLs). D1-D2 OLs, immature OLs; D4 OLs, mature OLs.

## Results

### BMP4 is expressed primarily in oligodendroglial lineage cells

BMP4 is a secreted protein and the mRNA expression better reflects its cellular origin. To determine cellular source of BMP4 *in vivo*, we purified different cell populations from the brain by MACS (**Fig. 1A** followed by mRNA quantification. We found that BMP4 was highly enriched in O4^+^ oligodendroglial lineage (**Fig. 1B**) which consists of late post-mitotic OPCs and immature OLs (Armstrong et al., 1992). To determine BMP4 dynamics within the oligodendroglial lineage, we employed primary OPC culture and differentiation system (Zhang et al., 2018b) and demonstrated that BMP4 was low in OPCs and rapidly upregulated (by >150-fold) in differentiating OLs (**Fig. 1C**), a dynamic pattern correlated with that of TCF7l2 (**Fig. 1D**) (Hammond et al., 2015), suggesting a potential link between BMP4 and TCF7l2 during OL differentiation.

### TCF7l2 deletion upregulates BMP4 and activates canonical BMP4 signaling *in vivo*

Our prior study reported an upregulation of BMP4 in the CNS of TCF7l2 cKO mutants (Hammond et al., 2015). To determine the regulation of BMP4 at the cellular level, we deleted TCF7l2 in the oligodendroglial lineage using time-conditioned Olig2-CreER^T2^ and assessed BMP4 expression *in vivo* 24 hours after tamoxifen treatment (**Fig. 2A**). During this short time window, TCF7l2 deletion-elicited OL differentiation defect (Hammond et al., 2015; Zhao et al., 2016) was not dramatically prominent, as indicated by ~28% decrease in the cell number of mRNA in situ hybridization (ISH) for the mature OL marker MAG (**Fig. 2B**). We found a significant increase of *Bmp4* mRNA in TCF7l2 cKO spinal cord demonstrated by RT-qPCR (**Fig. 2C**) and confirmed by ISH (**Fig. 2D**). Combined ISH and immunohistochemistry (IHC) showed that BMP4 was activated in Sox10^+^ cells in the spinal cord of TCF7l2 cKO mice (**Fig. 2E**), indicating that TCF7l2 represses oligodendroglial BMP4 expression *in vivo*.

**Fig. 2.**
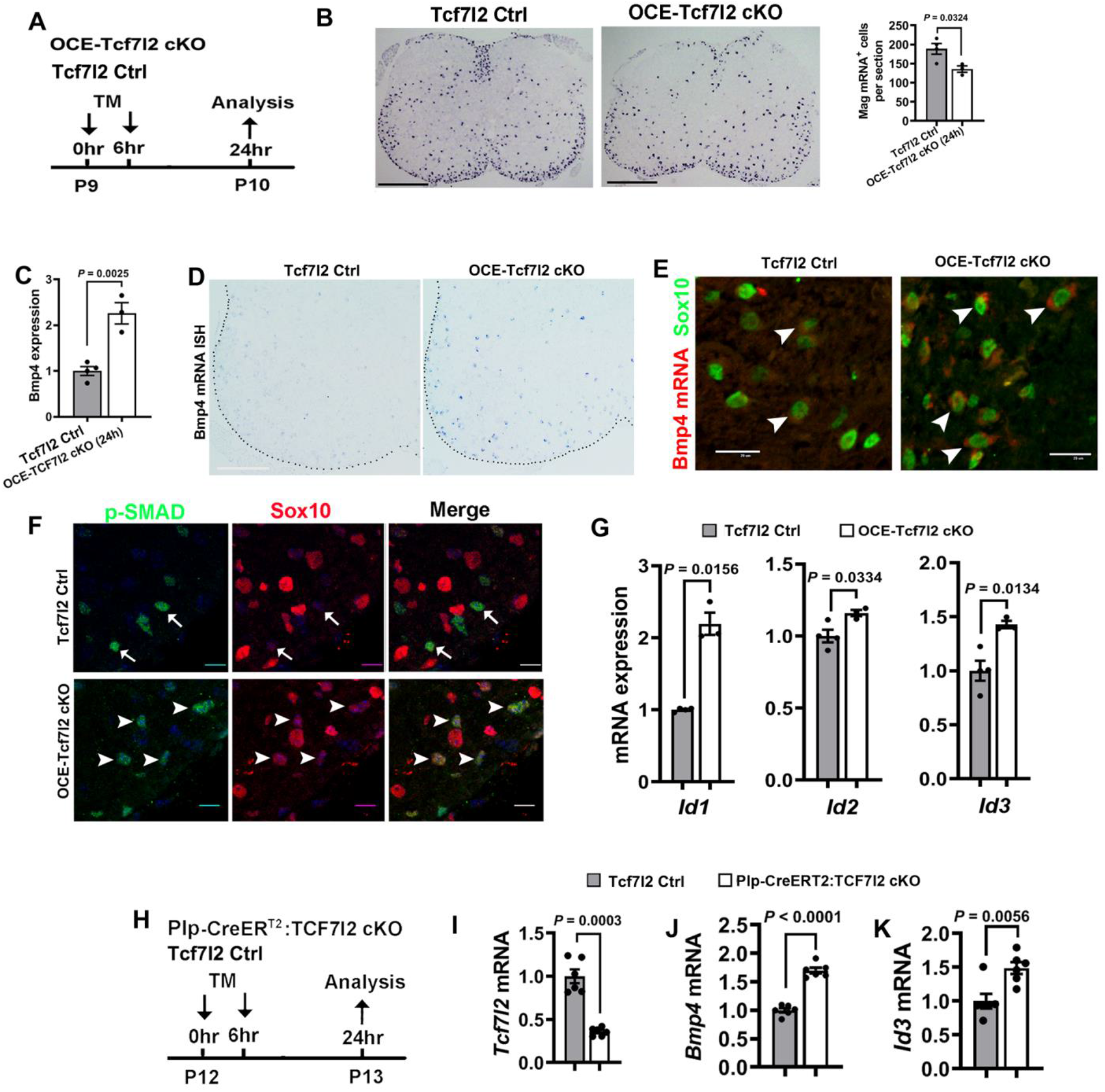
TCF7l2 deletion activates BMP4 and BMP4-mediated signaling in oligodendroglial lineage. A, experimental design for **B-G.** OCE-Tcf7l2 cKO (*Olig2-CreER^T2^*:*Tcf7l2*^fl/fl^ and Tcf7l2 Ctrl (*Tcf7l2*^fl/fl^, or *Tcf7l2*^fl/+^) mice were injected with tamoxifen at P9 and analyzed 24 hours later. **B**, In situ hybridization (ISH) and quantification of myelin associated glycoprotein (Mag) mRNA. **C**, RT-qPCR assay for *Bmp4* mRNA in the spinal cord. **D**, ISH showing *Bmp4* mRNA upregulation in the spinal cord. **E**, ISH of *Bmp4* mRNA and immunohistochemistry (IHC) of oligodendroglial marker Sox10 in the ventral white matter of spinal cord. **F**, double IHC of phosphorylated SMAD1/5 (p-SMAD) and Sox10 in the ventral white matter of spinal cord. Note that p-SMAD was barely detectable in Sox10^+^ cells of Tcf7l2 Ctrl spinal cord (arrows point to Sox10^-^/p-SMAD^+^ cells) while was activated in OCE-Tcf7l2 cKO spinal cord (arrowheads point to SOX10^+^/p-SMAD^+^ cells). **G**, *Id1, Id2*, and *Id3* expression quantified by RT-qPCR in the spinal cord. **H-K**, experimental design (**H**) and RT-qPCR quantification for *Tcf7l2* (**I**), *Bmp4* (**J**), and *Id3* (**K**) in the brain. Scale bars (in μm): **B**, 500; **D**, 200; **E**, 20; **F**, 10. Please see **Table 1** for statistical details throughout this study.

BMP4 activates canonical BMP signaling through phosphorylating SMAD1/5 (Retting et al., 2009), thus the phosphorylated form of SMAD1/5 (p-SMAD1/5) is a reliable surrogate for the signaling activation. We next used IHC antibodies specifically recognizing p-SMAD1/5 to assess BMP signaling activation at the cellular level. We found that p-SMAD1/5 was primarily observed in Sox10-negative cells, presumably astroglial lineage cells in Tcf7l2 Ctrl mice (**Fig. 2F**, arrows), suggesting that, under normal condition, the activity of BMP/SMAD signaling is minimal in oligodendroglial lineage cells. In contrast, p-SMAD1/5 immunoreactive signals were consistently observed in Sox10-expressing oligodendroglial cells (**Fig. 2F**, arrowheads) in TCF7l2 cKO mice. Furthermore, the expression of canonical BMP signaling target genes *Id1, Id2*, and *Id3* was significantly elevated in TCF7l2 cKO mice (**Fig. 2G**). These data suggest that TCF7l2 normally represses BMP4/SMAD signaling and its conditional depletion results in the activation of the signaling in oligodendroglial lineage cells. The upregulation of BMP4 and the signaling activity were confirmed in the brain by an independent *Plp1-CreER^T2^, Tcf7l2*^fl/fl^ mutant mice (**Fig. 2H-K**), suggesting that TCF7l2 controls BMP4 and the canonical BMP4 signaling activity in an autocrine manner not only in the spinal cord but also in the brain.

BMP4 enhances astrocyte generation at the expense of oligodendrogenesis (Gomes et al., 2003; Grinspan, 2020; Wu et al., 2012). To determine the effect of BMP4 upregulation elicited by TCF7l2 cKO on astrocyte development, we quantified astrocyte density *in vivo*. No significant difference was observed in the number of Sox9^+^BLBP^+^ or Sox9^+^S100β^+^ astrocytes (**Fig. 3A-D**), nor the expression of mature astrocyte markers GFAP and GLT-1 (**Fig. 3E-F**) between TCF7l2 Ctrl and cKO mice. Collectively, our data demonstrate that the Wnt effector TCF7l2 represses BMP4 expression and controls autocrine BMP signaling activity during OL differentiation.

**Figure 3.**
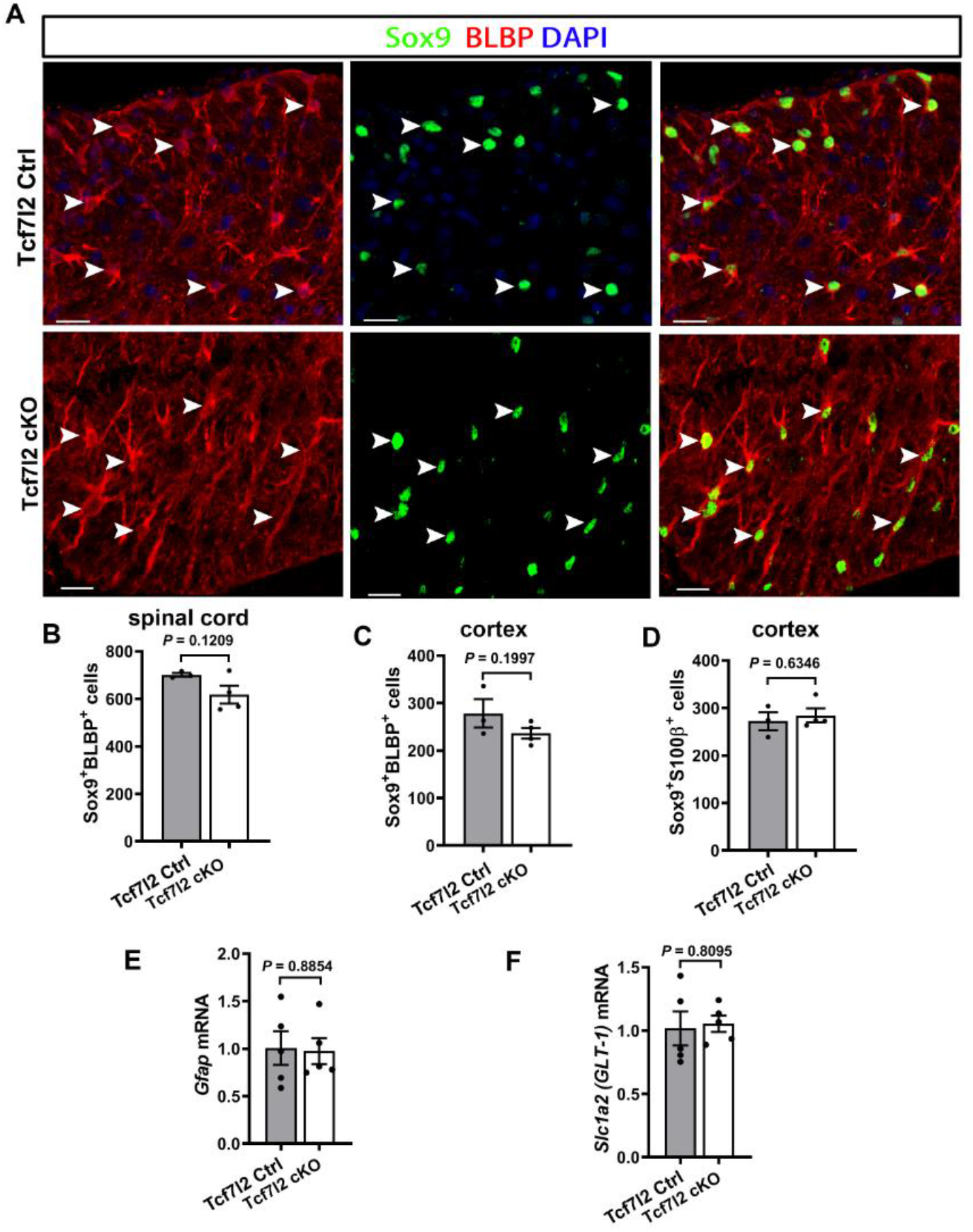
oligodendroglial TCF7l2 depletion does not affect astrocyte development in the CNS. *Olig2-CreER^T2^, Tcf7l2*^fl/fl^ (Tcf7l2 cKO) and TCF7l2 Ctrl mice (*Tcf7*^fl/fl^) were treated with tamoxifen at P6 and P7 and analyzed at P14. **A**, representative confocal images showing Sox9^+^BLBP^+^ astrocytes (arrowheads) in the spinal cord white matter. Scale bar=20μm. **B-C**, density (#/mm^2^) of Sox9^+^BLBP^+^ astrocytes in the spinal cord white matter (**B**) and forebrain cerebral cortex (**C**). **D**, Sox9^+^S100β^+^ astrocytes in the forebrain cerebral cortex. **E-F**, RT-qPCR quantification of mature astrocyte markers *Gfap* and *Slc1a2* in the spinal cord.

### TCF7l2 directly represses BMP4 transcription

A repressive role of TCF7l2 in BMP4 has not been reported in oligodendroglial lineage cells. Bioinformatic analysis showed that the intronic promoter of the mouse *Bmp4* gene (Thompson et al., 2003) contained a putative DNA binding motif of TCF7l2 (**Fig. 4A**). Chromatin immunoprecipitation (ChIP) followed by qPCR (ChIP-qPCR) confirmed the physical binding of TCF7l2 to this regulatory element (**Fig. 4B**). To determine the functional output of TCF7l2 binding, we sought to analyze the effect of TCF7l2 overexpression on the intronic *Bmp4* promoter-driven luciferase activity.

**Figure 4.**
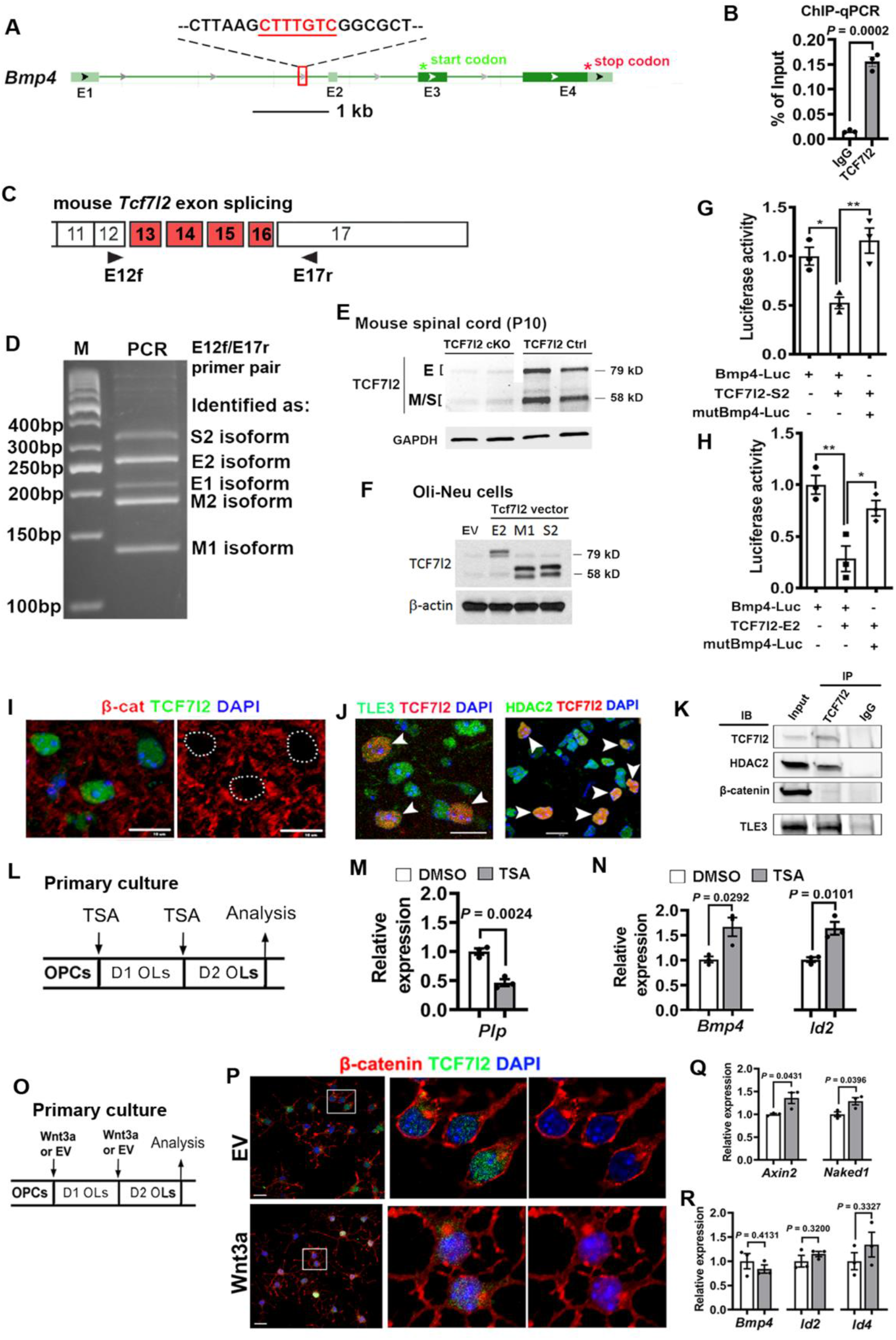
TCF7l2 directly represses BMP4 expression. A, putative TCF7l2-binding site located at ~317bp upstream of the exon 2 (E2) of the mouse *Bmp4* gene. **B**, ChIP-qPCR demonstrating TCF7l2 binding to the intronic *Bmp4* promoter shown in **A. C**, schematic diagram showing alternative splicing of the mouse *Tcf7l2* gene (Weise et al., 2010) and PCR primer pairs (E12f and E17r) used for amplifying mRNA transcripts of different splicing isoforms. **D**, PCR amplicons using the cDNA prepared from P10 mouse spinal cord. The amplicons were TA-cloned and sequenced. All three major isoforms (E, S, and M) were expressed in the spinal cord. **E**, Western blot assay for TCF7l2. Both ~79 kD (E isoform) and ~58 kD (M/S isoforms, similar molecular weight) bands were present in the mouse spinal cord and substantially downregulated in P10 TCF7l2 cKO spinal cord (*Olig2-CreER*^T2^:*Tcf7l2*^fl/fl^, tamoxifen injection at P9). TCF7l2 (clone C48H11, Cell Signaling #2569) was used. GAPDH serves as a loading control. *F*, Western blot showing vector-mediated expression of the representative mouse TCF7l2 isoforms (E2, M1, and S2) in the mouse oligodendroglial cell line Oli-Neu cells. Vectors were obtained from (Weise et al., 2010). EV, empty vector. **G-H**, luciferase assays for Bmp4-luc and mutated Bmp4-luc (mutBmp4) in the presence of TCF7l2 S2 (**G**) or E2 (**H**) isoforms. **I-J**, double IHC showing TCF7l2 is co-labeled with HDAC2 and co-repressor TLE3 (**J**, arrowheads) but not β-catenin (**I**, dashed circles) in P7 spinal cord. Scale bar=10μm. **K**, TCF7l2 Co-IP and Western blot detection of HDAC2, β-catenin, and TLE3 in P7 spinal cord. **L**, experimental design for **M-N**. Primary OPCs were purified from neonatal mouse brain. The HDAC inhibitor TSA (10ng/ml) or DMSO control was included in the differentiation medium for 48 hours (medium change at 24 hr) prior to analysis. The dose was chosen based on previous study (Marin-Husstege et al., 2002). **M**, RT-qPCR assay showing significant reduction of mature OL marker Plp in TSA-treated OLs. **N**, RT-qPCR assay for *Bmp4* and *Id2* in primary OLs treated with DMSO or TSA. **O**, experimental design for **P-R**. Primary mouse OPCs was transduced with empty control vector (EV) and vector expressing Wnt3a, a canonical Wnt ligand, at the time of differentiation and analyzed 48 hours after transduction. **P**, confocal images of single optical slice showing localization of β-catenin in Wnt3a-transduced and empty vector-transduced OLs. The boxed areas were shown at higher magnification on the right. Scale bar=10μm. **Q-R**, Relative mRNA levels of Wnt/β-catenin signaling targets *Axin2* and *Naked1* (**Q**) and *Bmp4* and BMP4 signaling targets *Id2* and *Id4*. White bars, Ctrl; gray bars: Wnt3a.

TCF7l2 have three major categories of alternative splicing isoforms (E, M, and S) depending the inclusion/exclusion of exons 13-16 (**Fig 4C**) (Weise et al., 2010). PCR amplification using primers targeting *Tcf7l2* exon12 and exon 17 in combination with DNA sequencing (**Fig. 4C**) showed that *Tcf7l2* mRNA transcripts corresponding to E, M, and S isoforms were present in the murine spinal cord (**Fig. 4D**). The expression of TCF7l2 E isoform (~79kD) and M/S isoforms (~58kD) was further confirmed by Western blot assay at the protein level in the spinal cord (**Fig. 4E**) and in primary brain OLs (Hammond et al., 2015). We therefore tested the effects of the representative E2 and S2 isoform expression (**Fig. 4F**) on *Bmp4* transcription activity. To this end, we cloned a 439-bp intronic promoter sequence (Thompson et al., 2003) into the luciferase reporter pGL2 vector and co-transduced with TCF7l2-E2 or S2 vectors into U87, a human primary glioblastoma cell line. Our results demonstrated that both E2 and S2 isoforms inhibited *Bmp4*-luciferase activity (**Fig. 4G-H**). Importantly, mutation of the TCF7l2 binding site abolished the repressive effect (**Fig. 4G-H**), suggesting that TCF7l2 represses BMP4 expression at the transcriptional level.

TCF7l2 harbors several functional domains including a β-catenin-binding domain, a Groucho/TLE-binding domain, and a high mobility group (HMG) box DNA-binding domain (Weise et al., 2010) which specifically recognizes CTTTG DNA motif on target genes (Hatzis et al., 2008). The gene regulation (activation or repression) by TCF7l2 is determined by its interacting co-activators or co-repressors in a cell type-dependent manner (Hurlstone and Clevers, 2002). To gain mechanistic insights into how TCF7l2 inhibits BMP4, we assessed the interaction of TCF7l2 with β-catenin and histone deacetylase 2 (HDAC2) *in vivo*, both of which have been shown to compete for TCF7l2 binding (Ye et al., 2009). TCF7l2 was histologically co-labeled with HDAC2 (**Fig. 4J**) but not β-catenin (**Fig. 4I**). Co-IP followed by Western blot demonstrated that TCF7l2 bound primarily to HDAC2 yet minimally to β-catenin (**Fig. 4K**). The binding of the co-activator β-catenin to TCF7l2 displaces the co-repressor Groucho/TLE from TCF7l2 (Clevers, 2006). The co-localization and interaction of TLE3 with TCF7l2 (**Fig. 4J, K**) also supported a minimal β-catenin binding to TCF7l2 under physiological conditions. To determine the involvement of HDAC2 in TCF7l2-mediated BMP4 repression, we treated primary OLs with Trichostatin A (TSA), a pan-HDAC inhibitor (**Fig. 4L**). In line with previous conclusion (Marin-Husstege et al., 2002; Ye et al., 2009), TSA remarkably decreased OL differentiation, evidenced by reduced expression of myelin protein gene *Plp* (**Fig. 4M**). Interestingly, *Bmp4* and canonical BMP4 signaling target *Id2* were significantly upregulated in TSA-treated OL culture (**Fig. 4N**), suggesting that the deacetylation activity of HDAC2 is required for TCF7l2-mediated BMP4 repression.

To determine whether “enforced” binding of β-catenin to TCF7l2 plays a role in TCF7l2-mediated BMP4 repression, we overexpressed Wnt3a, a typical canonical Wnt ligand, in primary OLs (**Fig. 4O**). In sharp contrast to cytoplasmic β-catenin in the control condition (**Fig. 4P,** upper panels), Wnt3a expression resulted in β-catenin localization in TCF7l2^+^DAPI^+^ nuclei (**Fig. 4P,** lower panels) and led to Wnt/β-catenin activation, as evidenced by the elevation in canonical Wnt targets *Axin2* and *Naked1* (**Fig. 4Q**). However, Wnt3a expression did not perturb the expression of BMP4 and canonical BMP/SMAD signaling targets *Id2* and *Id4* (**Fig. 4R**). Together, these data suggest that TCF7l2 inhibits BMP4 expression through recruiting the repressive co-factors HDAC2 and TLE3 to *Bmp4* cis-regulatory element and that β-catenin plays a dispensable role in TCF7l2-mediated BMP4 repression.

### TCF7l2 promotes OL differentiation by repressing BMP4 *in vitro*

To determine the biological significance of TCF7l2 in repressing BMP4, we overexpressed TCF7l2 in primary OLs and analyzed OL differentiation (**Fig. 5A-C**). TCF7l2 overexpression remarkably elevated the density of ramified MBP^+^Sox10^+^ OLs (**Fig. 5D**, left) by a greater than 2-fold (**Fig. 5D**, right) [% ramified Sox10^+^MBP^+^ cells: 8.34 ± 0.45% EV, 18.97± 2.80% TCF7l2-OE, t_(4)_ = 3.75, *P* = 0.0199] and significantly increased the expression of myelin protein genes (*Mbp, Plp*, and *Mog*), newly differentiated OL marker *Enpp6*, and pro-differentiation factor *Myrf* (**Fig. 5E**), indicating that TCF7l2 is sufficient for promoting OL differentiation in a cell autonomous manner. Consistent with the inhibitory role in BMP4 transcription, TCF7l2 overexpression reduced the amount of BMP4 protein secreted into the culture medium (**Fig. 5F**), decreased the number of Sox10^+^ oligodendrocytes that were positive for p-SMAD1/5 (**Fig. 5G**) [% p-SMAD^+^Sox10^+^ cells: 36.5 ± 1.1% EV, 28.9 ± 2.0% TCF7l2-OE, t_(4)_ = 3.298, *P* = 0.03], and reduced the expression of the canonical BMP4 signaling target *Id2* (**Fig. 5H**), suggesting that TCF7l2 promotes OL differentiation by inhibiting autocrine BMP4 and dampens BMP4-mediated canonical BMP/SMAD signaling activity. We did not find significant differences in the expression of canonical Wnt target genes (Guo et al., 2015) in between EV and TCF7l2-OE groups (**Fig. 5I**), indicating that TCF7l2 plays a minor role in transcriptionally activating Wnt/β-catenin signaling in oligodendroglial lineage cells (Hammond et al., 2015). BMP4 treatment (Fig. **5J**) elevated the canonical BMP/SMAD signaling activity in primary OLs as evidenced by Id2 upregulation upon BMP4 treatment (**Fig. 5K**). We found that BMP4 treatment abolished the upregulation of *Plp* expression elicited by TCF7l2 overexpression (**Fig. 5L**). Collectively, these data suggest that TCF7l2 promotes OL differentiation through controlling autocrine BMP4-mediated signaling.

**Figure 5.**
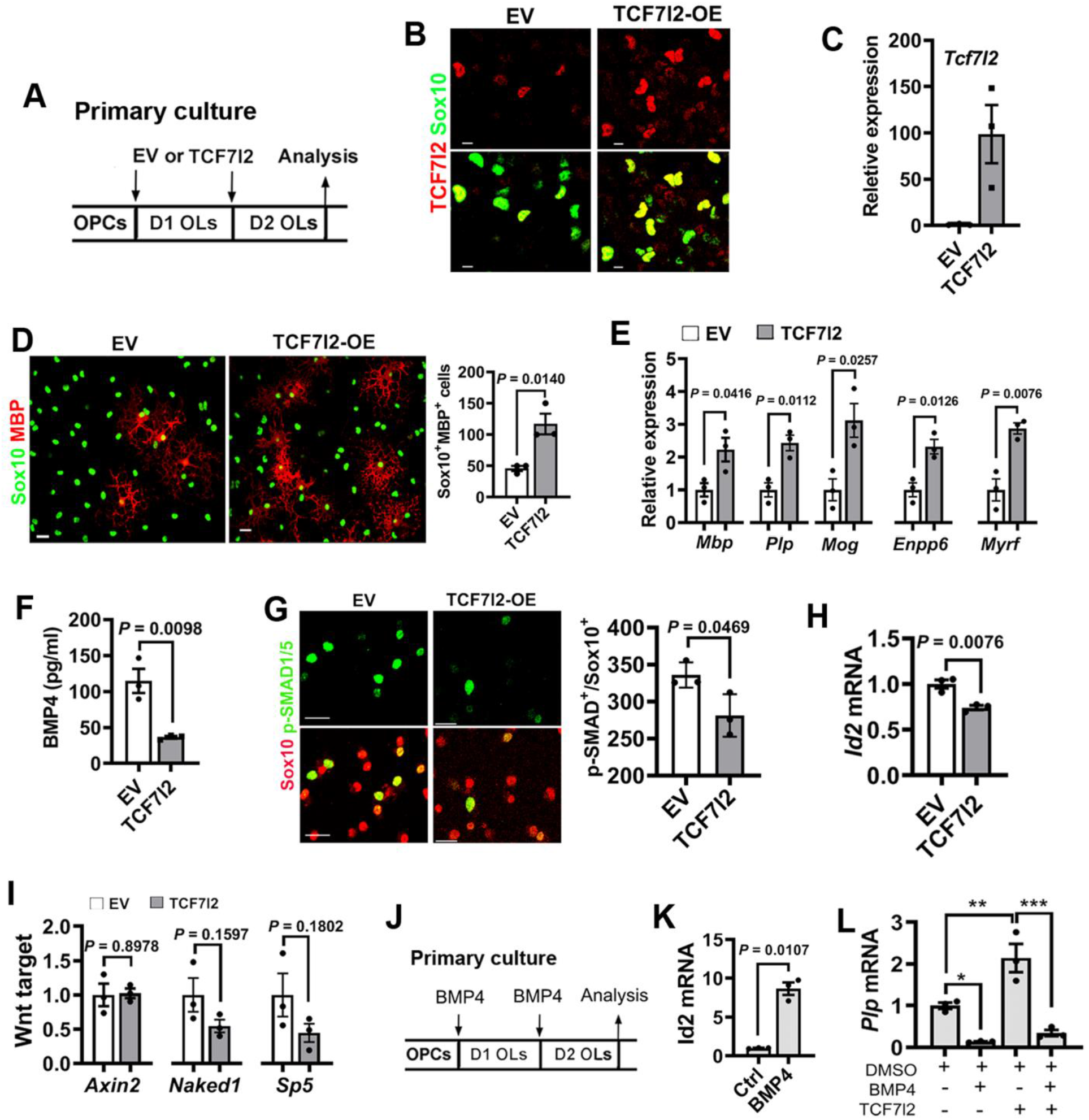
TCF7l2 overexpression represses BMP4-mediated signaling and promotes oligodendrocyte differentiation *in vitro*. **A**, experimental design for **B-I**. Primary OLs were transduced with empty vector (EV) or S2 isoform of TCF7l2 plasmid (TCF7l2 overexpression, TCF7l2-OE), and analyzed at 48 hours after transduction. **B-C**, representative confocal images of TCF7l2 and Sox10 (**B**) and RT-qPCR assay of *Tcf7l2* mRNA (**C**) in EV and TCF7l2-OE groups. **D**, immunocytochemistry (ICC) and quantification of Sox10^+^MBP^+^ cells transduced with EV or TCF7l2-OE vector. **E**, RT-qPCR quantification for myelin genes, newly formed OL marker *Enpp6*, and pro-differentiation factor *Myrf*. **F**, ELISA assay for BMP4 concentration in culture medium. **G**, representative confocal images and density of p-SMAD1/5^+^Sox10^+^ cells (#/mm^2^). **H**, relative mRNA levels of BMP4 signaling target *Id2* quantified by RT-qPCR. **I**, relative expression of canonical Wnt targets quantified by RT-qPCR. **J**, experimental design for **K-L**. BMP4 (10ng/ml) or DMSO control was included in the differentiation medium for 48 hours (medium change at 24 hr) prior to analysis. **K**, RT-qPCR quantification of canonical BMP4 signaling target Id2. Two-tailed t test, Welch’s corrected *t*_(2.009)_ = 9.529. **L**, RT-qPCR assay for OL marker *Plp* in primary OLs treated with BMP4, TCF7l2-OE, or both. Scale bars=10 μm in **B, D, G**.

### Disrupting oligodendroglial BMP4 rescues OL differentiation defects in TCF7l2 cKO mutants

To determine the physiological significance of TCF7l2-mediated BMP4 repression on OL differentiation *in vivo*, we generated *Olig2-CreER*^T2^, *Tcf7l2*^fl/fl^ (TCF7l2 single cKO), *Olig2-CreER*^T2^, *Tcf7l2*^fl/fl^, *Bmp4*^fl/fl^ (TCF7l2/BMP4 double cKO), and control mice. In line with the above observation (**Fig. 2**), TCF7l2 disruption (**Fig. 6A-B**) caused BMP4 upregulation (**Fig. 6C**) and BMP4/SMAD signaling activation (**Fig. 6D-E**) in Sox10^+^ oligodendroglial lineage cells (**Fig. 6F-G**) of TCF7l2 single cKO mutants. Simultaneous *Bmp4* deletion (**Fig. 6C**) normalized BMP/SMAD signaling activity in TCF7l2/BMP4 double cKO mice to the level in control mice (**Fig. 6D-G**).

**Fig. 6.**
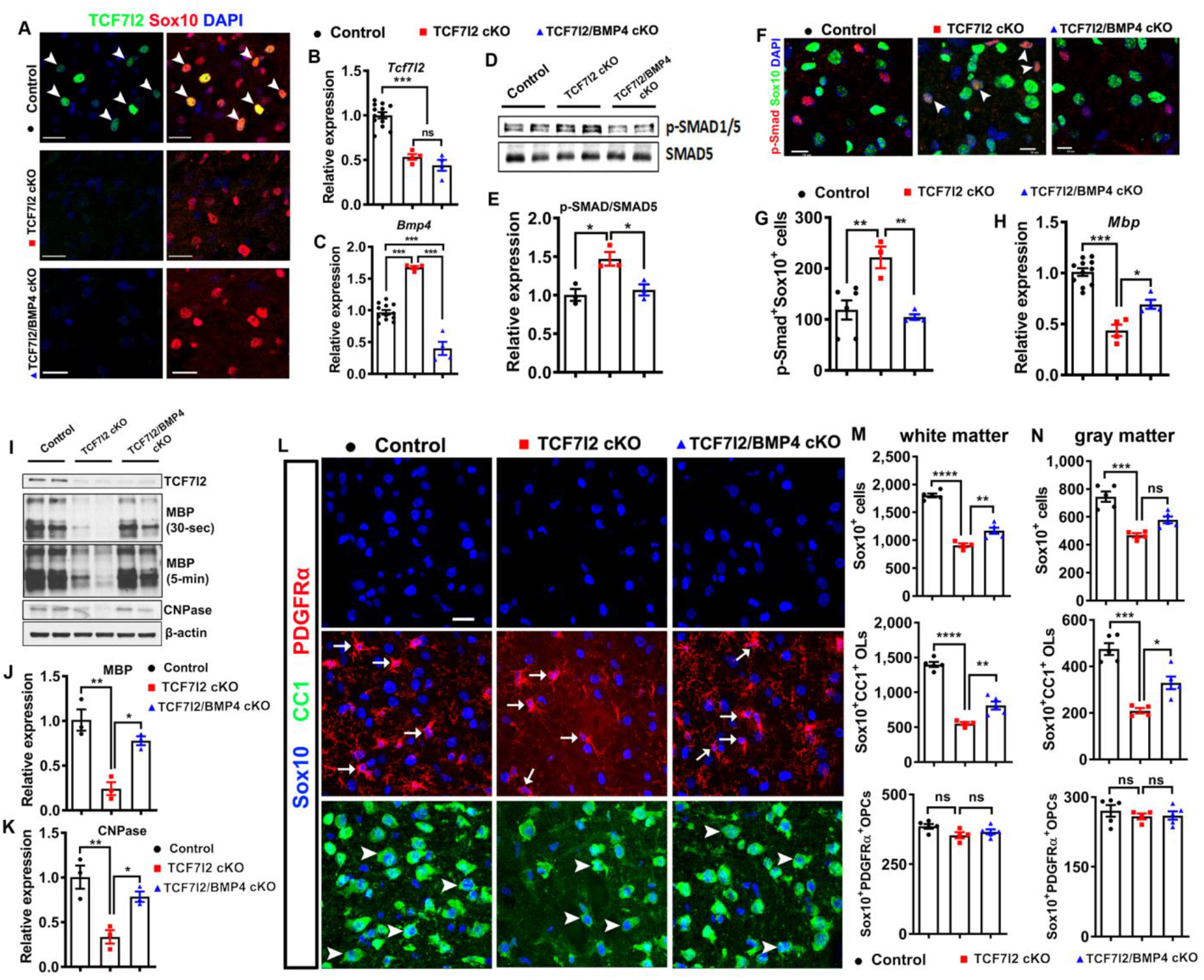
Disrupting BMP4 rescues TCF7l2 deletion-elicited inhibition of oligodendrocyte differentiation *in vivo*. *Olig2-CreER*^T2^, *Tcf7l2*^fl/fl^ (TCF7l2 cKO), *Olig2-CreER*^T2^, *Tcf7l2*^fl/fl^, *Bmp4*^fl/fl^ (TCF7l2/BMP4 cKO), and control mice were treated with tamoxifen at P6 and P7 and analyzed at P13. **A**, representative confocal images of TCF7l2 and Sox10 co-labeling cells (arrowheads) in the spinal cord. **B-C**, relative expression of *Tcf7l2* and *Bmp4* in the spinal cord assessed by RT-qPCR. **D-E**, representative Western images and quantification of p-SMAD1/5 over SMAD5 in the forebrain. **F-G**, representative confocal images and quantification of p-SMAD1/5 and Sox10 IHC (#/mm^2^) in the spinal cord white matter. **H**, relative *Mbp* expression quantified by RT-qPCR. **I-K**, representative Western blot images and quantification of TCF7l2, MBP, and CNPase in the forebrain. β-actin serves as an internal loading control. **L**, representative confocal images of Sox10, CC1, and PDGFRα IHC in the spinal cord white matter. Arrowheads, Sox10^+^CC1^+^ OLs; arrows, Sox10^+^PDGFRα^+^ OPCs. **M-N**, densities (#/mm^2^) of total Sox10^+^ oligodendroglial lineage cells, Sox10^+^CC1^+^ OLs, and Sox10^+^PDGFRα^+^ OPCs in the white matter (**M**) and gray matter (**N**) of spinal cord. Scale bars, 20 μm in A, 10 μm in F and L.

Consistent with previous reports (Hammond et al., 2015; Zhao et al., 2016), TCF7l2 deficiency remarkably decreased the expression of myelin basic protein MBP and CNPase at the mRNA (**Fig. 6H**) and protein level (**Fig. 6I-K**) and significantly diminished OL differentiation (**Fig. 6L-N**) in TCF7l2 single cKO mice compared with control mice. Simultaneous BMP4 deletion remarkably alleviated the defects in myelin gene expression (**Fig. 6H-K**) and OL differentiation (**Fig. 6L-N**) in TCF7l2/BMP4 double cKO compared with TCF7l2 single cKO mice. Thus, our genetic results convincingly demonstrate that BMP4 dysregulation is a downstream molecular mechanism underlying OL differentiation defects elicited by TCF7l2 cKO *in vivo*.

To determine the effect of BMP4 depletion alone on OL differentiation *in vivo*, we analyzed *Pdgfra-CreER*^T2^, *Bmp4*^fl/fl^ (BMP4 cKO) mice and age-matched control mice. Despite a greater than 4-fold BMP4 reduction in BMP4 cKO mice compared with BMP4 Ctrl littermates (**Fig. 7A**), we did not find significant differences in the number of differentiated OLs and OPCs (**Fig. 7B-D**), myelin gene expression (**Fig. 7E**), or the rate of OL differentiation indicated by the number of APC^+^TCF7l2^+^ (Zhang et al., 2018a) newly generated OLs (**Fig. 7F-I**). Despite >4 fold reduction in BMP4, BMP4 cKO mice maintained a comparatively normal level of BMP/SMAD signaling activity, as assessed by similar levels of canonical BMP/SMAD target genes *Id1* and *Id2* between BMP4 Ctrl and cKO mice (Fig. 7J). These data suggest that other BMP ligands, for example BMP2 and BMP6 from other cell types (See et al., 2007), may be redundant with BMP4 in maintaining normal BMP/SMAD signaling activity and OL differentiation in BMP4 cKO mice.

**Figure 7.**
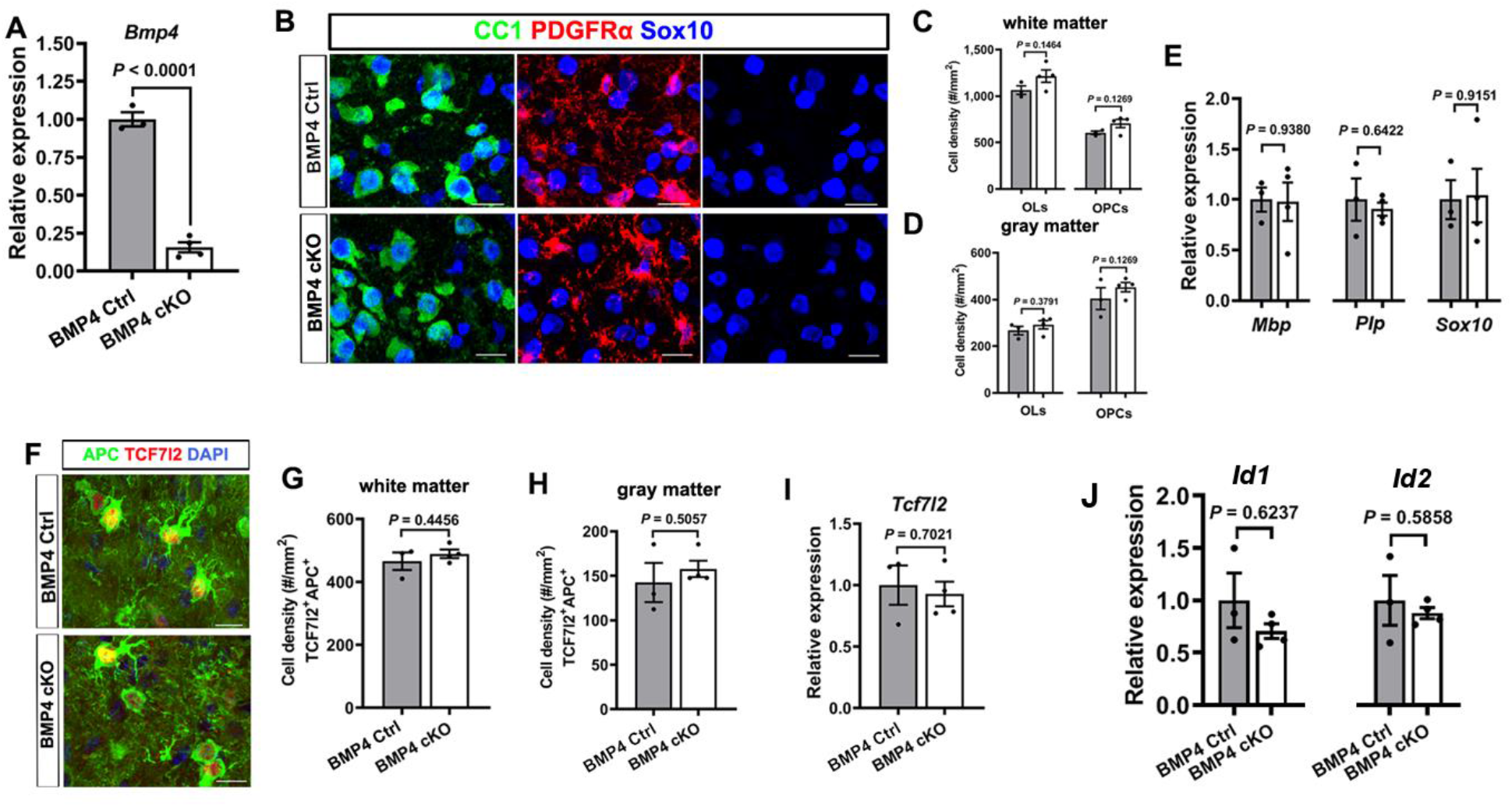
BMP4 disruption alone does not affect oligodendrocyte differentiation in vivo. *Pdgfrα-CreER*^T2^, *Bmp4*^fl/fl^ (BMP4 cKO) and BMP4 Ctrl (*Bmp4*^fl/fl^) mice were treated with tamoxifen at P5 and P6 and the spinal cord was analyzed at P10. **A**, relative *Bmp4* expression by RT-qPCR. **B-D**, representative images of Sox10/CC1/PDGFRα IHC (**B**) and densities of Sox10^+^CC1^+^ OLs and Sox10^+^PDGFRα^+^ OPCs in the spinal cord white (**C**) and gray (**D**) matter. Scale bar=10μm. **E**, relative expression of myelin protein gene *Mbp* and *Plp* and pan-oligodendroglial marker *Sox10* quantified by RT-qPCR. **F-H**, representative confocal image of newly differentiated OLs labeled by APC/TCF7l2 (**F**), and quantification of APC^+^/TCF7l2^+^ cells in the white (**G**) and gray (**H**) matter. Scale bar=10μm. **I**, relative Tcf7l2 expression in the spinal cord by RT-qPCR. **J**, relative expression of canonical BMP signaling targets *Id1* and *Id2* quantified by RT-qPCR.

## Discussion

By employing *in vitro* primary culture and genetic mouse models in combination with biochemical and molecular analyses, we made several novel observations: 1) oligodendroglial lineage is the primary cellular source of BMP4 in the CNS; 2) TCF7l2 controls the level of autocrine BMP4 signaling activity in oligodendroglial lineage cells by transcriptionally repressing *Bmp4* gene; and 3) TCF7l2 promotes OL differentiation by dampening autocrine BMP4-mediatd signaling *in vitro* and *in vivo*.

Our previous studies have shown that TCF7l2 is transiently upregulated in newly-formed OLs (Hammond et al., 2015) that are positive for adenomatous polyposis coli (APC), a Wnt/β-catenin negative regulator (Lang et al., 2013) (**Fig. 8A**). Based on our findings, we proposed a “Wnt-independent and BMP4-dependent” role of TCF7l2 in promoting OL differentiation (**Fig. 8B**). In this working model, APC limits the nuclear availability of β-catenin for TCF7l2 binding and Wnt/β-catenin activation whereas TCF7l2 binds to the cis-regulatory elements of *Bmp4* genes and directly represses *Bmp4* transcription likely by recruiting co-repressors HDAC2 and TLE3, both of which lack DNA-binding domains. Disrupting the DNA-binding TCF7l2 leads to uncontrolled BMP4 expression and inhibits OL differentiation through BMP4-mediated autocrine signaling activation.

**Fig. 8.**
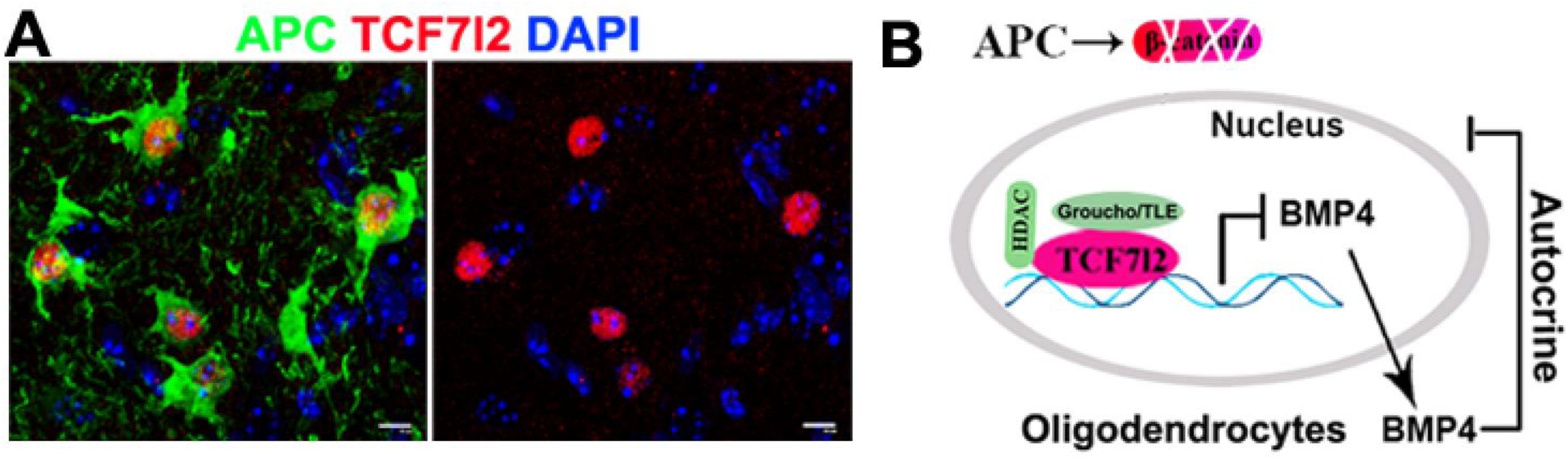
Conceptual framework of BMP4 repression by TCF7l2 in promoting oligodendrocyte differentiation. **A**, All TCF7l2^+^ oligodendroglial cells are positive for APC, a negative regulator of Wnt/β-catenin signaling. **B**, proposed working model of “Wnt-independent and BMP4-dependent” role of TCF7l2 in promoting OL differentiation.

Previous data show that Wnt/β-catenin signaling regulates the timely differentiation of OPCs into OLs (Dai et al., 2014). Our working model of Wnt-independent function of TCF7l2 does not conflict with the conclusion by Dai and colleagues (Dai et al., 2014). The TCF/LEF family consists of TCF7, TCF7l1, TCF7l2, and LEF1 all of which could transcriptionally mediate Wnt/β-catenin signaling. It is possible that that other TCF/LEF family member may mediate the effect of Wnt/ β-catenin signaling on oligodendroglial differentiation. Consistent with this possibility, our previous data (Hammond et al., 2015) demonstrated that Wnt/β-catenin signaling activation elicited by APC disruption (Lang et al., 2013) remarkably upregulates the expression of the Wnt effector LEF1 in oligodendroglial lineage cells. We are current employing oligodendroglial-specific APC/LEF1 double cKO system to prove or falsify our hypothesis.

TCF7l2 is upregulated primarily in active MS lesions (with extensive remyelination) but minimally in chronic MS lesions (with scarce remyelination) (Fancy et al., 2009). Recent studies demonstrate an elevation of BMP4 in both active and chronic MS lesions (Costa et al., 2019; Harnisch et al., 2019) although the cell types producing BMP4 need to be defined (Grinspan, 2020). The repressive TCF7l2-BMP4 regulation suggests that augmenting TCF7l2 may have synergistic potential in dampening BMP4 signaling, which is dysregulated in chronic MS lesions (Grinspan, 2015, 2020) and in promoting OL differentiation and remyelination given the multi-modal actions of TCF7l2 in propelling OL lineage progression and maturation (Zhao et al., 2016).

The proposed working model (**Fig. 8B**) may also be applicable to a broader cell lineage. Beyond its role in CNS oligodendroglial lineage, the Wnt effector TCF7l2 regulates a spectrum of biological processes of other lineage cells, such as hepatic metabolism (Boj et al., 2012), colorectal tumor genesis (Angus-Hill et al., 2011; Wenzel et al., 2020), adipocyte function (Chen et al., 2018), and heart development (Ye et al., 2019) where BMP signaling plays crucial roles (Brazil et al., 2015; Wang et al., 2014). The connection between TCF7l2 and autocrine BMP4 signaling has not been reported in these cell types. In this regard, our take-home message (Fig. 8B) point to a new direction for studying the crosstalk between TCF7l2 and BMP4/SMAD signaling pathways in regulating the development and function of TCF7l2-expressing and BMP/Wnt-operative cells in the body. Supporting this claim, recent data suggest that TCF7l2 overexpression downregulates *Bmp4* mRNA in a cardiac cell line (Ye et al., 2019).

In summary, our *in vitro* and *in vivo* data unravel a previously unrecognized role of the Wnt effector TCF7l2 in promoting OL differentiation by negatively controlling BMP4-mediated autocrine signaling. Given that BMP4 is dysregulated in chronic MS lesions, our findings of the biological function of TCF7l2-BMP4 connection in OL differentiation indicate that enforced TCF7l2 expression may be a potential therapeutic option in promoting OL and myelin regeneration in chronic MS lesions and animal mode.

## Acknowledgments

We thank Dr. Andreas Hecht (University of Freiburg, Germany) for providing TCF7l2-E2, M1, and S2 expression vectors and Dr. Gino Cortopassi (UC Davis) for providing U87 cells. This study was funded by NIH/NINDS (R21NS109790 and R01NS094559) and by Shriners Hospitals for Children (85107-NCA-19, 84553-NCA-18, and 84307-NCAL).

